# Host PIK3C3 promotes *Shigella flexneri* spread from cell to cell through vacuole formation

**DOI:** 10.1101/2024.10.31.621244

**Authors:** Steven J Rolland, Zachary J Lifschin, Erin A Weddle, Lauren K Yum, Tsuyoshi Miyake, Daniel A Engel, Hervé F Agaisse

**Author notes:** Corresponding author: Hervé F Agaisse.

## Abstract

*Shigella flexneri* is a human intracellular pathogen responsible for bacillary dysentery (bloody diarrhea). *S. flexneri* invades colonic epithelial cells and spreads from cell to cell, leading to massive epithelial cell fenestration, a critical determinant of pathogenesis. Cell-to-cell spread relies on actin-based motility, which leads to formation of membrane protrusions, as bacteria project into adjacent cells. Membrane protrusions resolve into intermediate structures termed vacuole-like protrusions (VLPs), which remain attached to the primary infected cell by a membranous tether. The resolution of the membranous tether leads to formation of double-membrane vacuoles (DMVs), from which *S. flexneri* escapes to gain access to the cytosol of adjacent cells. Here, we identify the class III PI3K family member PIK3C3 as a critical determinant of *S. flexneri* cell-to-cell spread. Inhibition of PIK3C3 decreased the size of infection foci formed by *S. flexneri* in HT-29 cells. Tracking experiments using live-fluorescence confocal microscopy showed that PIK3C3 is required for efficient resolution of VLPs into DMVs. PIK3C3-dependent accumulation of PtdIns(3)P at the VLP membrane in adjacent cells correlated with the transient recruitment of the membrane scission machinery component Dynamin 2 at the neck of VLPs at the time of DMV formation. By contrast, *Listeria monocytogenes* did not form VLPs and protrusions resolved directly into DMVs. However, PIK3C3 was also required for *L. monocytogenes* dissemination, but at the stage of vacuole escape. Finally, we showed that PIK3C3 inhibition decreased *S. flexneri* dissemination in the infant rabbit model of shigellosis. We propose a model of *Shigella* dissemination in which vacuole formation relies on the PIK3C3-dependent accumulation of PtdIns(3)P at the VLP stage of cell-to-cell spread, thereby supporting the resolution of VLPs into DMVs through recruitment of the membrane scission machinery component, DNM2.

**Author summary:** *Shigella flexneri* is an intracellular pathogen causing bacillary dysentery, a disease responsible for more than 200,000 deaths each year in the world. With the lack of efficient vaccines and the dramatic increase in multi-drug-resistant clinical isolates, it is critical to better understand *Shigella* pathogenesis to suggest new therapeutic treatments. Previous studies demonstrated that invasion of epithelial cells in the human colon and subsequent spread from cell to cell are critical determinants of pathogenesis. Cell-to-cell spread relies on manipulation of the host cell actin cytoskeleton supporting bacterial movement in the cytosol of infected cells. At cell-cell contacts, bacteria project into adjacent cells through formation of membrane protrusions that resolve into vacuoles from which the bacteria escape to gain access to the cytosol of adjacent cells. Here, we show that the host cell kinase PIK3C3 is critical for efficient cell-to-cell spread through resolution of protrusions into vacuoles. Our work suggests that inhibitors of PIK3C3 may represent novel avenues of therapeutic intervention in shigellosis.

## Introduction

The human pathogen *Shigella flexneri* is the causative agent of bacillary dysentery, or shigellosis. The disease is the second-leading cause of diarrheal diseases with 270 million cases including more than 200,000 deaths per year worldwide (Khalil et al., 2018; Kotloff et al., 2018). Shigellosis is characterized by invasion and dissemination of *S. flexneri* into the colonic epithelium, leading to epithelial fenestration, massive infiltration of immune cells and bloody diarrhea (Mathan and Mathan, 1991; Yum et al., 2019). The emergence and isolation of multi-drug-resistant species worldwide, and the lack of efficient vaccines emphasize the importance of better understanding *S. flexneri* pathogenesis of (Kotloff *et al*., 2018; Puzari et al., 2017; Thorley et al., 2023).

The capacity of *S. flexneri* to invade colonic epithelial cells and spread from cell to cell is critical for pathogenesis and relies on the expression specific virulence factors, including the type three secretion system (T3SS) and its associated bacterial effectors (Agaisse, 2016; Sansonetti, 1991; Weddle and Agaisse, 2018a). Once *S. flexneri* escapes the primary vacuole and reaches the host cytoplasm, the polar expression of the bacterial protein IcsA recruits host cell Neural Wiskott–Aldrich syndrome protein (N-WASP, also known as WASL), which recruits and activates the actin-related protein 2/3 complex (ARP2/3), leading to actin-based motility in the cytosol of infected cells (Egile et al., 1999; Suzuki et al., 1998; Welch and Way, 2013). At cell-cell contacts, motile bacteria project into adjacent cells through formation of membrane protrusions. The engulfment of membrane protrusions by adjacent cells leads to the formation of double membrane vacuoles from which the pathogen escapes to gain access to the cytosolic compartment of adjacent cells, thereby by achieving cell-to-cell spread (Agaisse, 2016; Weddle and Agaisse, 2018a).

Numerous mechanisms are required for successful formation of membrane protrusions. *S. flexneri* disseminates efficiently in polarized epithelial cells that not only establish E-cadherin-mediated cell-cell contacts (Sansonetti et al., 1994), but also STK11-dependent lateral junctions competent for tyrosine kinase signaling (Dragoi and Agaisse, 2014a; b). In addition, *S. flexneri* requires the presence of tricellulin for efficient plaque formation and appears to preferentially form protrusions at tricellular junctions (Fukumatsu et al., 2012). The formation of protrusions relies on the forces generated by the actin cytoskeleton in order to deform the plasma membrane and protrude into adjacent cells. Actin network formation in protrusions is not only supported by the ARP2/3 complex, but also by specific cytoskeleton factors, including the formins mDia1/2 (Heindl et al., 2010) and myosin X (Bishai et al., 2013). Protrusion formation necessitates the release of membrane tensions mediated through interaction of the T3SS translocase IpaC with the junctional protein B-catenin (Duncan-Lowey et al., 2020). Proper elongation of protrusions also relies on local exocytosis potentially contributing to the release of membrane tension or providing required membrane material at the site of protrusion formation (Herath et al., 2021). In addition to membrane tension at cell-cell contacts, *S. flexneri* must overcome *de novo* actin polymerization occurring underneath the inner membrane in protrusions. This is achieved through secretion of the T3SS effector protein IpgD, which hydrolyses PIP2, a signaling event that presumably prevents the recruitment of the yet uncharacterized *de novo* actin polymerization machinery (Koseoglu et al., 2022).

The resolution of protrusions into vacuoles is probably the least understood aspect of *S. flexneri* dissemination (Weddle and Agaisse, 2018a). After a brief phase of protrusion elongation, the collapse of the actin network in the neck of the formed protrusions leads to the generation of intermediate structures termed vacuole-like protrusions (VLPs), which remain attached to the primary infected cell by a membranous tether (Dragoi and Agaisse, 2015). This step requires several cellular signaling events, including PIK3C2α-dependent formation of PtdIns(3)P in the inner membrane of protrusions and T3SS-dependent activation of tyrosine kinase signaling (Dragoi and Agaisse, 2014a; 2015; Kuehl et al., 2014). The collapse of the actin cytoskeleton in the protrusion neck is thought to be a mandatory step for subsequent remodeling of the membrane tether of VLPs, resulting in vacuole formation (Dragoi and Agaisse, 2015).

In addition to membrane and cytoskeleton related events occurring in the primary infected cell, the neighboring cells play an active role in the engulfment of protrusions. It was first proposed that the engulfment of protrusions involves non-canonical clathrin-mediated endocytosis dependent on PI3Ks signaling (Fukumatsu *et al*., 2012). Moreover, a recent study reported the recruitment of caveolin-1, cavin-2, actin and VASP to protrusions in adjacent cells, suggesting that *S. flexneri* might also rely on the caveolin endocytic pathway and actin polymerization for efficient protrusion engulfment (Dhanda and Guttman, 2023). Importantly, chemical inhibition or genetic depletion of the membrane scission machinery DNM2 associated with endocytic events have been shown to affect *S. flexneri* plaque formation and therefore cell-to-cell spread (Fukumatsu *et al*., 2012; Lum et al., 2013). However, the exact role of DNM2 in *S. flexneri* dissemination remains unclear.

Here, we found that inhibitors of phosphatidylinositol 3-kinase class 3 (PIK3C3) decrease the efficiency *S. flexneri* cell-to-cell spread. We used time lapse microscopy to demonstrate that PIK3C3-dependent formation of PtdIns(3)P at the outer membrane of protrusions is required for efficient resolution of VLPs into double membrane vacuoles (DMVs). PtdIns(3)P formation was required for the recruitment of the membrane scission machinery DNM2. DNM2 was transiently recruited to the neck of VLPs at the bacterial pole, which coincided with the time of the VLP membranous tether scission and DMV formation. Finally, we show that inhibiting PIK3C3 attenuated *S. flexneri* spread *in vivo* in the infant rabbit model of shigellosis.

## Results

### PIK3C3 promotes *Shigella flexneri* dissemination in HT-29 cells

In a chemical screen for host cell kinases required for *S. flexneri* dissemination, we identified seven potent compounds, including the class III phosphatidylinositol 3- phosphate kinase (PIK3C3) inhibitor VPS34-IN1 (S1 Table), that decreased the size of the infection foci formed by *S. flexneri* in HT-29 cells. Confirming the specific activity of VPS34-IN1, we observed a VPS34-IN1 concentration-dependent decrease of the size of the infection foci formed in HT-29 colonic epithelial infected with CFP-expressing *S. flexneri* (Fig 1A, 1D). We confirmed these results with SAR405, another specific and potent inhibitor of PIK3C3 (Ronan et al., 2014). As previously observed with VPS34-IN1, SAR405 treatment decreased *S. flexneri* dissemination in a concentration-dependent manner (Fig 1B, 1E). Finally, we conducted RNA interference experiments with two PIK3C3-specific siRNA duplexes and confirmed that efficient PIK3C3 depletion (Fig 1G, 1H) decreased *S. flexneri* dissemination (Fig 1C, 1F). We next tested whether PIK3C3 inhibition could negatively impact actin-based motility. To this end, we infected HT-29 cells in presence of VPS34-IN1 and determined the percentage of bacteria associated with an actin tail using confocal microscopy. The approach revealed that VPS34-IN1 did not significantly affect the percentage of bacteria with an actin tail, suggesting that PIK3C3 inhibition does not impact *S. flexneri* dissemination through actin-based motility (Fig 1I). Altogether, these chemical and genetic experiments suggest that the PI3K family member PIK3C3 promotes *S. flexneri* cell-to cell-spread in colonic epithelial cells.

**Fig 1.**
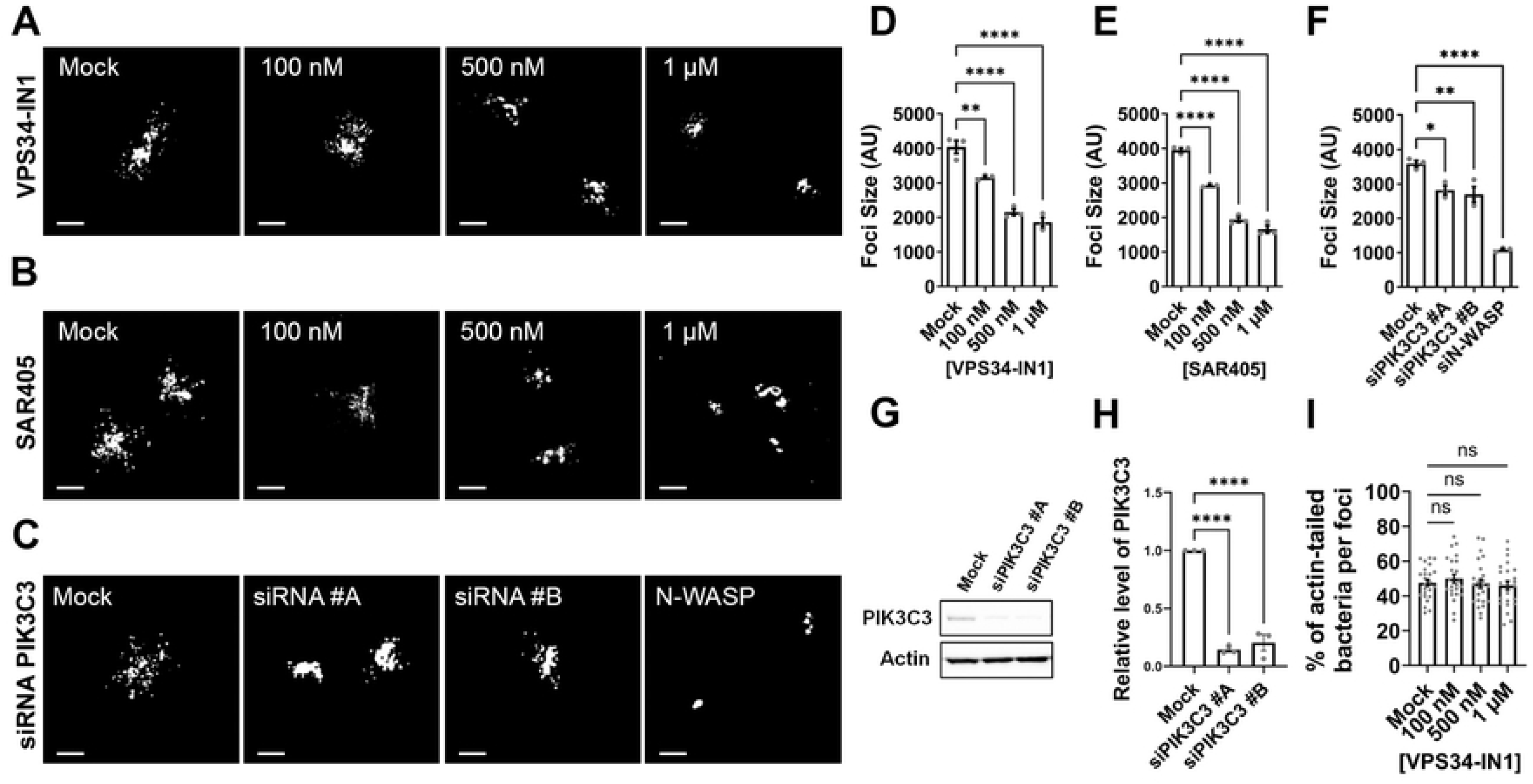
PIK3C3 promotes *Shigella flexneri* cell-to-cell spread in HT-29 cells. **(A-C)** Representative images showing infection foci formed by CFP-expressing *S. flexneri* in HT-29 cells 8 hours post-infection in presence of the specific PIK3C3 inhibitor VPS34-IN1 (A) or SAR405 (B), or upon treatment with siRNA duplexes targeting PIK3C3 or N-WASP (as positive control for no spread) (C). Scale bar, 50 μm. **(D-F)** Quantification of foci size (area) in arbitrary units 8 hours post-infection in presence of VPS34-IN1 (D), or SAR405 (E) or upon treatment with siRNA duplexes targeting PIK3C3 (F). Each dot represents the average of one biological replicate. At least 50 infection foci were analyzed in each of three independent biological replicates. Error bars represent standard error of the mean. Statistical analysis, one-way ANOVA with Dunnett’s multiple comparisons test; ns, not significant; *, p<0.05; **, p<0.005; ****, p<0.0001. **(G)** Western blot showing knockdown efficiency of two siRNA duplexes targeting PIK3C3. **(H)** Quantification of the knockdown efficiency as shown in (G) for three biological replicates. PIK3C3 was normalized to loading control (Actin) and knockdown efficiency was calculated relative to mock treated cells. Error bars represent standard error of the mean. Statistical analysis, one-way ANOVA with Dunnett’s multiple comparisons test; ns, not significant; ****, p<0.0001. **(I)** Percentage of bacteria displaying an actin tail per infection focus 4 hours post-infection in presence of VPS34-IN1. Eight foci were analyzed per replicate and three biological replicates were performed. Error bars represent standard error of the mean. Statistical analysis, one-way ANOVA with Dunnett’s multiple comparisons test; ns, not significant.

### PIK3C3 promotes *S. flexneri* cell-to-cell spread through the resolution of VLPs into DMVs

Previous studies showed that *S. flexneri* spread from cell to cell is a multi-step process, including protrusion formation, transition into VLPs, resolution of VLPs into DMVs, and DMV escape (Fig. 2A) (Dragoi and Agaisse, 2015). To determine which step was affected by PIK3C3 inhibition, we used live-imaging confocal microscopy to track the formation of protrusions, VLPs and DMVs in real-time during *S. flexneri* cell-to-cell spread (Dragoi and Agaisse, 2015; Koseoglu *et al*., 2022; Weddle and Agaisse, 2018b; Weddle et al., 2022). In mock conditions, 49.7% of bacteria went through all stages of cell-to-cell spread and successfully spread into adjacent cells after 4h30 min (Fig 2B-C, S1 Fig, red). In stark contrast, only 6.3% of bacteria successfully spread from cell to cell when treated with the PIK3C3 inhibitor. Analyzing the failure situations, we found that an equal percentage of bacteria failed to escape DMVs in mock- and VPS34-IN1-treated cells (Fig 2B-C, S1 Fig, yellow). In mock-treated samples, 23% of bacteria formed protrusions failed to form VLPs and retracted to primary infected cells (Fig 2B-C, S1 Fig, dark blue). In VPS34-IN1-treated cells however, bacteria formed protrusions that either failed to form VLPs and retracted to the primary infected cells (Fig 2B, light blue followed by dark blue and Fig 2C, dark blue, 43.2%) or form VLPs that failed to resolve into DMVs, and then retracted to primary cells (Fig 2B, light blue followed by purple and then dark blue, and Fig 2C, purple, 25.3%). As previously reported (Dragoi and Agaisse, 2015), VLP formation represented a strict commitment to DMV formation in mock-treated cells (Fig 2B, purple (VLP) systematically followed by yellow (DMV)). Remarkably, in VPS34-IN1-treated cells, 25.2% of bacteria failed in VLPs that did not commit to DMV formation (Fig 2C, purple; S2 Fig). Finally, we found that the time spent in VLPs and DMVs, but not the time spent in protrusions, was significantly increased in the presence of VPS34-IN1 (Fig 2D-2G). Altogether, these results reveal a critical role for PIK3C3 in the resolution of VLPs into DMVs during *S. flexneri* dissemination.

**Fig 2.**
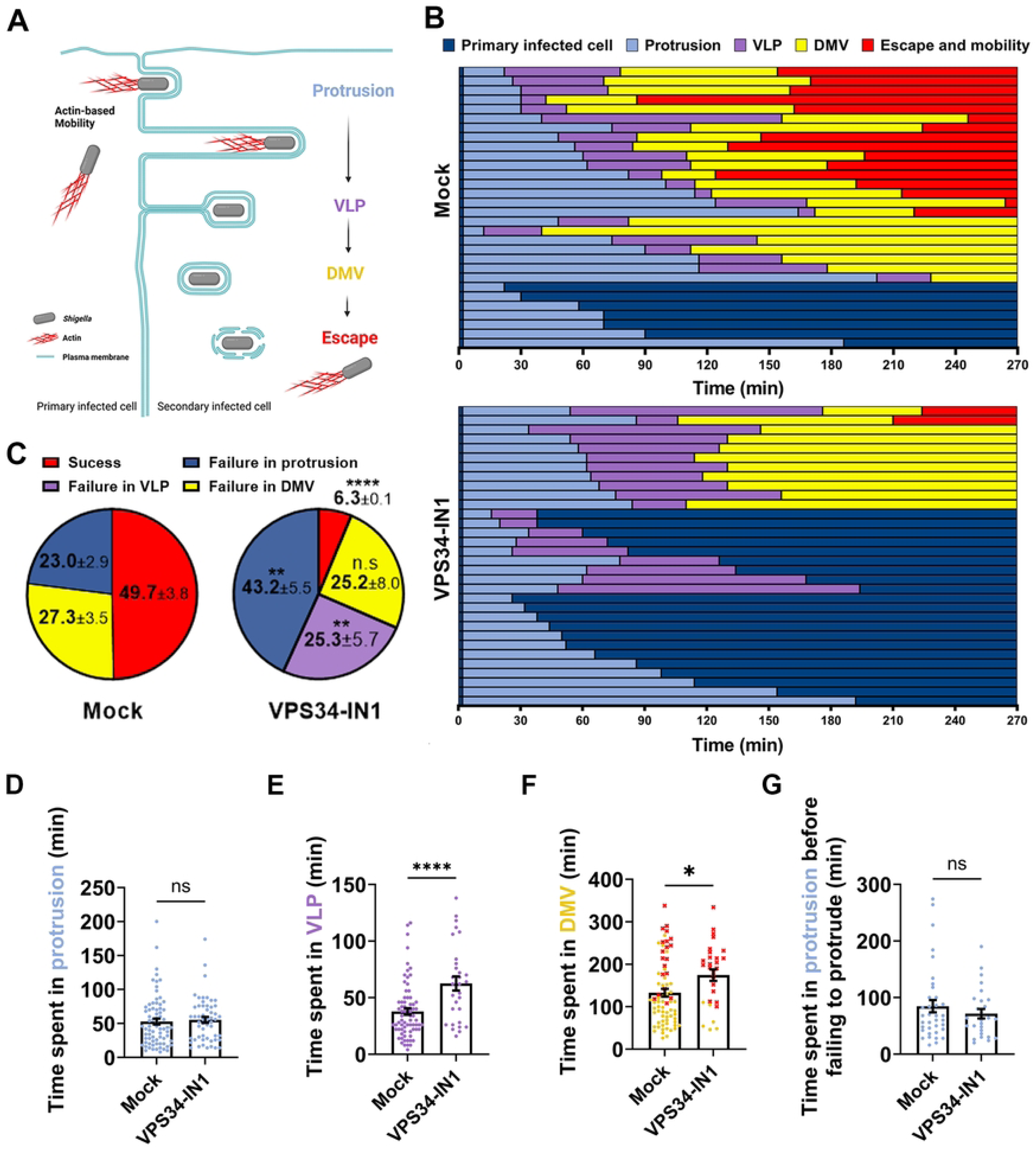
PIK3C3 promotes *Shigella flexneri* cell-to-cell spread through resolution of VLPs into DMVs. **(A)** Schematic representation of *S. flexneri* cell-to-cell spread. The bacterium (grey) displays actin-based motility (red filaments) and projects into adjacent cells at cell-cell contacts through formation of membrane protrusions (light blue). The collapse of the actin cytoskeleton in the protrusion neck results in the formation of an intermediate structure termed Vacuole-Like Protrusions (VLPs, purple). The resolution of the membrane tether linking VLPs to primarily infected cells lead to the formation of Double Membrane Vacuoles (DMVs, yellow). DMV escape grants the bacteria access to the cytosol of secondary infected cell where they resume actin-based motility (red). **(B)** Representative tracking analysis using live-fluorescence confocal microscopy of HT-29 cells infected with CFP-expressing *S. flexneri* in absence (left) or presence (right) of VPS34-IN1 at 500 nM. Each bar represents the tracking of a single bacterium over 4h30. At least 30 bacteria were tracked for each condition per biological replicate. Color code (bottom right): dark blue, primarily infected cells; light blue, protrusion; purple, vacuole- like protrusion (VLP); yellow, double membrane vacuole (DMV) and red, escape and actin-based motility in the cytosol of adjacent cells. **(C)** Pie chart showing the proportion of bacteria failing at a given stage in three biological replicates. Means and standard error of the means are indicated. Statistical analysis, two-way ANOVA with Sidak’s multiple comparisons; ns, not significant; *, p<0.05; ***, p<0.001; ****, p<0.0001. **(D-F)** Time spent in protrusions (D), VLPs (E), or DMVs (F) for bacteria that successfully transitioned to the next stage of the dissemination process. Red dots correspond to bacteria that did not escape DMVs by the end of tracking. Error bars represent the standard error of the means. Statistical analysis, unpaired t-tests; ns, not significant; *, p<0.05; ****, p<0.0001. **(G)** Time spent in protrusions for all bacteria that failed to spread from cell-to-cell and did not form VLPs. Error bars represent the standard error of the means. Unpaired t-tests were performed, ns, not significant.

### PIK3C3 promotes *Listeria monocytogenes* cell-to-cell spread through DMV escape

Although displaying similar strategies of actin-based motility in the cytosol, *Listeria monocytogenes* and *S. flexneri* have evolved different strategies of cell-to-cell spread (Kuehl et al., 2015). As presented in Figure 2A, *S. flexneri* relies on VLP formation which results from the collapse of the actin cytoskeleton in protrusions (Dragoi and Agaisse, 2015). By contrast, *L. monocytogenes* relies on the critical role of the AIP1-dependent actin recycling machinery in fueling efficient actin assembly at the bacterial pole in protrusions (Talman et al., 2014). Vacuole formation results from the snapping of protrusion membrane, presumably as a consequence of the tension exerted on the membrane in stationary protrusions displaying continuous actin polymerization at the bacterial pole (Talman *et al*., 2014). Thus, the mechanisms leading to vacuole formation are different in *L. monocytogenes* and *S. flexneri* dissemination. To determine whether PIK3C3 plays a role in *L. monocytogenes* cell-to-cell spread, we infected HT-29 cells with GFP-expressing *L. monocytogenes* in presence or absence of PIK3C3 inhibitors. Since VLP formation does not occur during *L. monocytogenes* dissemination, we were surprised to observe a decrease in foci size in cells treated with VPS34-IN1 or SAR405 (Fig 3A-D).

**Fig 3.**
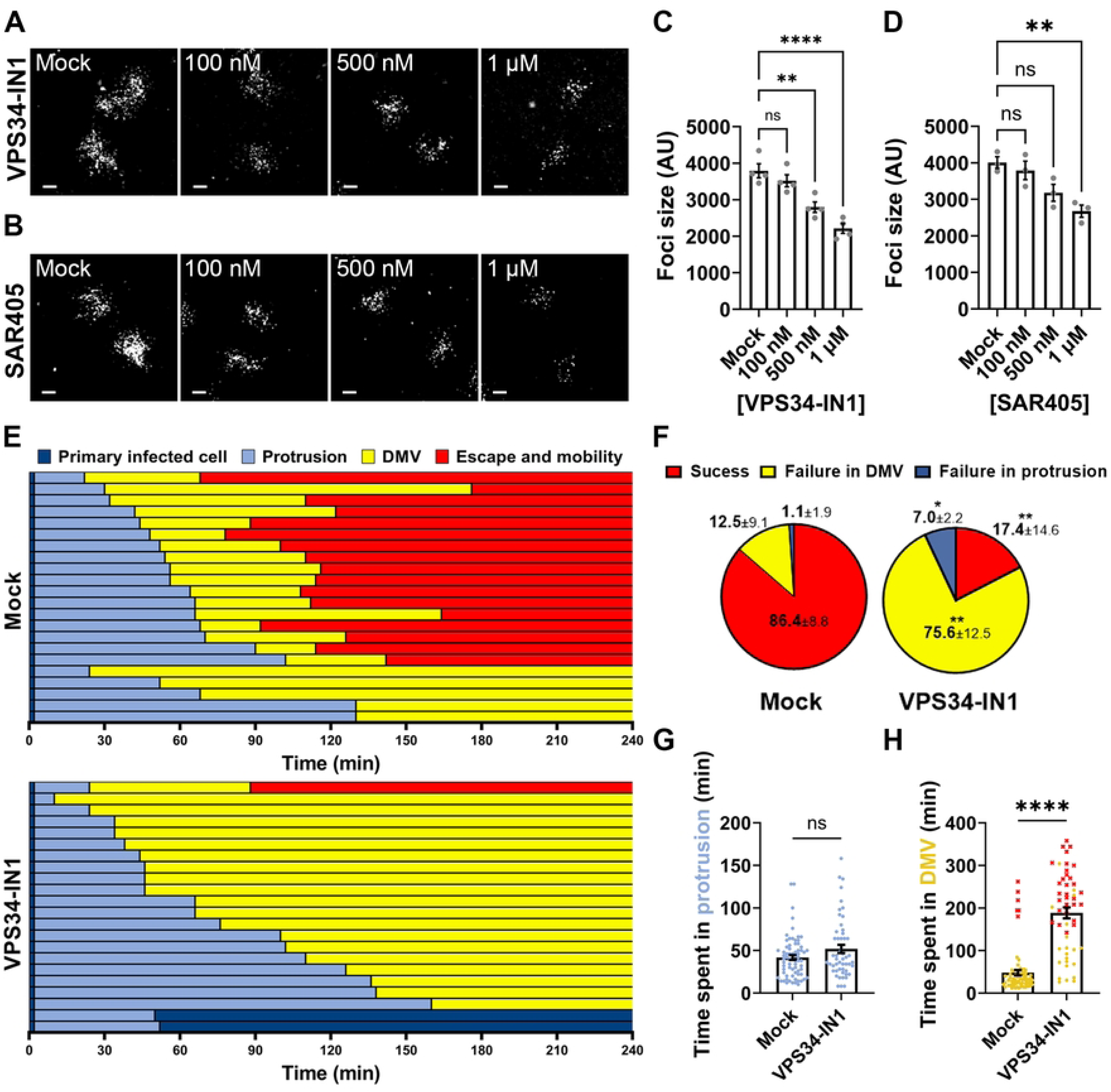
PIK3C3 promotes *Listeria monocytogenes* cell-to-cell spread in HT-29 cells through DMV escape. **(A-B)** Representative images showing infection foci formed 8 hours post-infection in HT-29 cells infected with GFP-expressing *L. monocytogenes* in presence of VPS34-IN1 (A) or SAR405 (B). Scale bar, 50 μm. **(C-D)** Quantification of foci size (area) in arbitrary units at 8 h post-infection in presence of VPS34-IN1 (C) or SAR405 (D). Each dot represents the average of one biological replicate. Three independent biological replicates were performed, each containing at least 50 foci. Error bars represent the standard error of the mean. Statistical analysis, one-way ANOVA with Dunnett’s multiple comparisons test; ns, not significant; **, p<0.005; ****, p<0.0001. **(E)** Representative tracking analysis using live-fluorescence confocal microscopy of HT-29 cells infected with CFP-expressing *L. monocytogenes* in absence (top panel) or presence of VPS34-IN1 at 500 nM (bottom). Each bar represents the tracking of a single bacterium over 4 hours. Color code (bottom right): dark blue, primarily infected cells; light blue, protrusion; yellow, double membrane vacuole (DMV) and red, escape and mobility in the cytosol of adjacent cells. **(F)** Pie chart showing the proportion of bacteria failing at a given stage in three biological replicates. Means and standard error of the means are indicated. Statistical analysis, two-way ANOVA with Sidak’s multiple comparisons; ns, not significant; *, p<0.05; ***, p<0.001; ****, p<0.0001. **(G-H)** Time spent in protrusions (G), or DMVs (H) for bacteria that transition to the next stage of the dissemination process. Each dot represents the time spent of a single bacterium in the corresponding phase. Red dots correspond to bacteria that did not escape DMVs by the end of tracking. Error bars represent the standard error of the means. Statistical analysis, unpaired t-tests; ns, not significant; *, p<0.05; ****, p<0.0001.

We next tracked *L. monocytogenes* cell-to-cell spread using live-imaging confocal microscopy to determine which dissemination step may be affected by PIK3C3 inhibition. As expected, the VLP intermediate formed during *S. flexneri* dissemination was not observed during *L. monocytogenes* cell-to-cell spread (S3 Fig). Tracking results showed that, while 86.4% of bacteria successfully spread from cell to cell in mock-treated cells, only 17.4% of bacteria gained access to the cytosol of adjacent cells in presence of PIK3C3 inhibitor (Fig 3E-F, S4 Fig, red). Strikingly, while the vast majority of bacteria successfully escaped DMVs in mock-treated cells, 75.6% of bacteria were trapped in DMVs in the presence of VPS34-IN1 (Fig 3E-F, S4 Fig). Additionally, among bacteria that successfully spread from cell to cell, PIK3C3 inhibition did not affect the time spent in protrusions, but the time spent in DMV was significantly increased (Fig 3G-H). Altogether, these results suggest a role for PIK3C3 in DMV escape during *L. monocytogenes* dissemination.

### PIK3C3 supports PtdIns(3)P formation during the VLP-to-DMV transition

PIK3C3 supports the production of phosphatidylinositol 3-phosphate (PtdIns(3)P) through the phosphorylation of the 3’OH group of phosphatidylinositol. PtdIns(3)P acts as a signaling molecule controlling the recruitment to various membrane-bound compartments of proteins displaying PtdIns(3)P-binding domains, such as the PX or FYVE domains. To determine whether PIK3C3 may lead to PtdIns(3)P production during the VLP-to-DMV transition, we conducted infection experiments using live-imaging confocal microscopy using a mixed population of mb-YFP-expressing HT-29 cells expressing or not an mCherry-PX probe. mCherry-PX consists of the fusion between the mCherry fluorescent protein and the PX domain of the p40^phox^ subunit of the superoxide-producing phagocyte NADPH-oxidase (or NCF4), which specifically binds PtdIns(3)P (Dragoi and Agaisse, 2015; Ellson et al., 2001; Kanai et al., 2001). Through imaging of bacteria spreading from a mCherry-PX non-expressing cell into a mCherry-PX expressing cell, we determined that the probe was transiently recruited to the plasma membrane surrounding bacteria in adjacent cells (Fig 4A). Tracking results revealed that PtdIns(3)P was produced just before or during the VLP stage and was always present during the VLP-to-DMV transition (Fig 4B, mock, purple followed by yellow, black bars). PtdIns(3)P was still present shortly after DMV formation, and then disappeared (Fig 4B, mock, yellow, black bars). In stark contrast with the mock-treated conditions, we did not observe the recruitment of the probe at any step of *S. flexneri* dissemination in presence of VPS34-IN1. We note that few bacteria successfully spread from cell to cell without formation of PtdIns(3)P, suggesting either short-lived PtdIns(3)P formation missed under our experimental procedure, or the existence of non-PIK3C3-dependent mechanisms of VLP-to-DMV transition. To confirm these visual observations, we quantified the levels of PX-mCherry probe fluorescence at membranes surrounding bacteria during cell-to-cell spread (Fig 4C). These results showed a significant 3-fold increase in PtdIns(3)P production over background signal at the time of VLP formation (Fig 4D-E) and no production in examples of failed protrusion- to-VLP transition (Fig 4F) or failed VLP-to-DMV transition (Fig 4G). We also tracked PtdIns(3)P formation during *L. monocytogenes* cell-to-cell spread. As expected given the role of PIK3C3 in DMV escape, PtdIns(3)P was formed to *L. monocytogenes*-containing DMVs after their formation, but not before (S3 Fig). Altogether, these results show that PIK3C3 supports PtdIns(3)P formation during the VLP-to-DMV transition and vacuole escape in cells infected with *S. flexneri* and *L. monocytogenes*, respectively.

**Fig 4.**
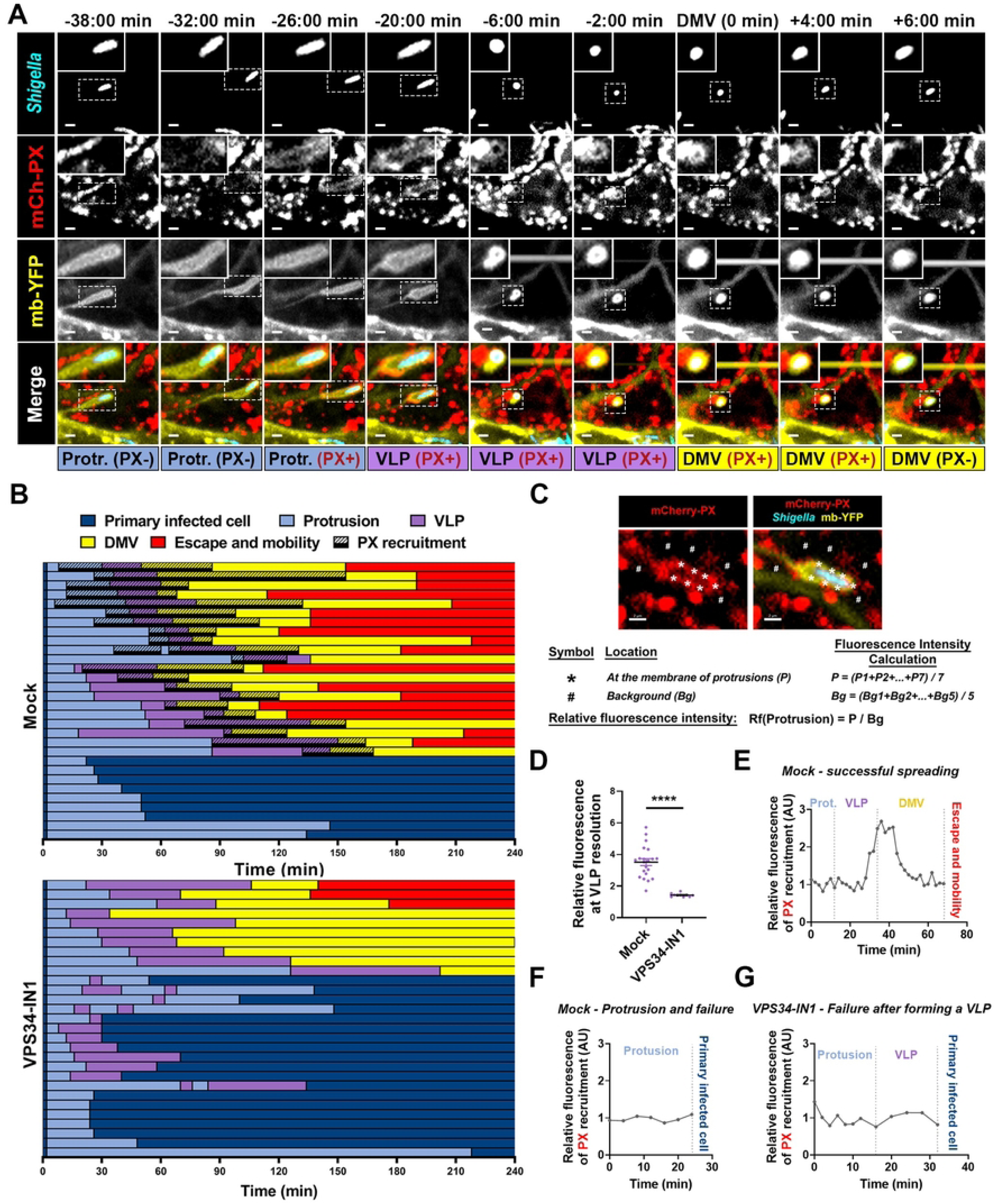
PIK3C3 promotes the resolution of VLPs into DMVs through formation of PtdIns(3)P at the protrusion/VLP membrane. **(A)** Representative images of membrane YFP-expressing cells infected with CFP-expressing *S. flexneri* dissemination from a PX- mCherry non-expressing cell to a PX-mCherry expressing cell. PX-mCherry specifically binds PtdIns(3)P. Scale bar, 2 μm. **(B)** Representative tracking analysis in HT-29 cells infected with CFP-expressing *S. flexneri* in absence (top panel) or presence of VPS34- IN1 at 500 nM (bottom panel). Each bar represents the tracking of a single bacterium spreading from a non-expressing cell into a PX-mCherry expressing cell over 4 hours. At least 30 bacteria were tracked for each condition. Color code: dark blue, primarily infected cells; light blue, protrusions; purple, vacuole-like protrusions (VLPs); yellow, double membrane vacuoles (DMVs) and red, cytosol of adjacent cells; black bar, recruitment of PX-mCherry. **(C)** Representative example of relative fluorescence intensity calculation. **(D)** Relative fluorescence of mCherry-PX at the VLP membrane the time frame before DMV formation. Error bars indicate standard deviation of the mean. Statistical analysis, unpaired t-tests were performed; ns, not significant; ****, p<0.0001. **(E)** Graphs representing the relative fluorescence intensity at the membrane of protrusion/VLP/DMV for an example of successful cell-to-cell spread. **(F-G)** Graphs representing the relative fluorescence intensity at the membrane of protrusions and VLPs for an example of failure in protrusion (F), and failure in VLP (G), in presence of VPS34-IN1(500 nM).

### DNM2 promotes *S. flexneri* cell-to-cell spread through VLP resolution

The VLP-to-DMV transition requires the resolution of the VLP membranous tether, suggesting the involvement of factors supporting membrane scission. Previously, chemical (Dynasore) and genetic (shRNA) approaches have suggested a role for DNM2 in *S. flexneri* dissemination (Fukumatsu *et al*., 2012; Lum *et al*., 2013). Moreover, DNM2 was observed around the protrusions formed in MK2 and Caco-2 cells using antibody staining and DNM2 fused to green fluorescent protein, respectively (Fukumatsu *et al*., 2012). However, a recent study showed a recruitment of DNM2 at the neck of protrusions formed by *S. flexneri* before detachment from the rest of host membrane (Plum et al., 2024), but not during protrusion formation as previously described (Fukumatsu *et al*., 2012). To address this apparent controversy about the recruitment of DNM2 and clarify its role during dissemination, we conducted infection experiments in HT-29 cells treated with DNM2-targeting siRNA duplexes. The efficiency of DNM2 knockdown was confirmed and quantified by western blot experiments (Fig S5C-D). DNM2 depletion led to a significant decrease in the size of the foci formed during *S. flexneri* dissemination (Fig S5A-B). This result confirms that DNM2 promotes *S. flexneri* cell-to-cell spread in a monolayer of HT-29 cells.

To visualize the recruitment of DNM2 during cell-to-cell spread, we tracked CFP-expressing *S. flexneri* spreading from mb-YFP HT-29 cells to mbYFP HT-29 cells expressing DNM2 fused to the mCherry fluorescent protein (DNM2-mCherry). We observed a signal corresponding to the DNM2-mCherry probe at the neck of VLPs, before their resolution into vacuoles, which disappeared once the DMV was formed (Fig 5A). Tracking results showed that successful formation of DMVs systematically correlated with recruitment of DNM2-mCherry at the neck of VLPs, just before the resolution of VLPs into DMVs (Fig 5B, mock, back bars). In the presence of the PIK3C3 inhibitor VPS34-IN1, bacteria forming DMVs showed DNM2 recruitment to VLPs. However, we did not observe significant DNM2 recruitment to VLPs that failed to transition into DMVs in VPS34-IN1 treated cells (Fig 5B). As opposed to the previous observations reported by Fukumatsu and collaborators (Fukumatsu *et al*., 2012), we did not observe any recruitment of DNM2 to protrusions. To confirm our observations, we quantified the fluorescence levels corresponding to the DNM2-mCherry probe at the neck of VLPs (Fig 5C, white circles) and at the membrane of protrusions (Fig 5C, white stars) normalized to background levels (Fig 5C, white sharp symbols). Tracking results showed a 2-fold increase in fluorescence levels in VLPs that successfully transitioned into DMVs in mock-treated cells (Fig 5D). However, fluorescence levels stayed low throughout the VLP stage and did not increase in cells treated with VPS34-IN1 (Fig 5E). Altogether these results expand on previous observations involving DNM2 in *S. flexneri* dissemination and suggest that the DNM2- mediated membrane scission machinery promotes cell-to-cell spread through resolution of VLPs into DMVs.

**Fig 5.**
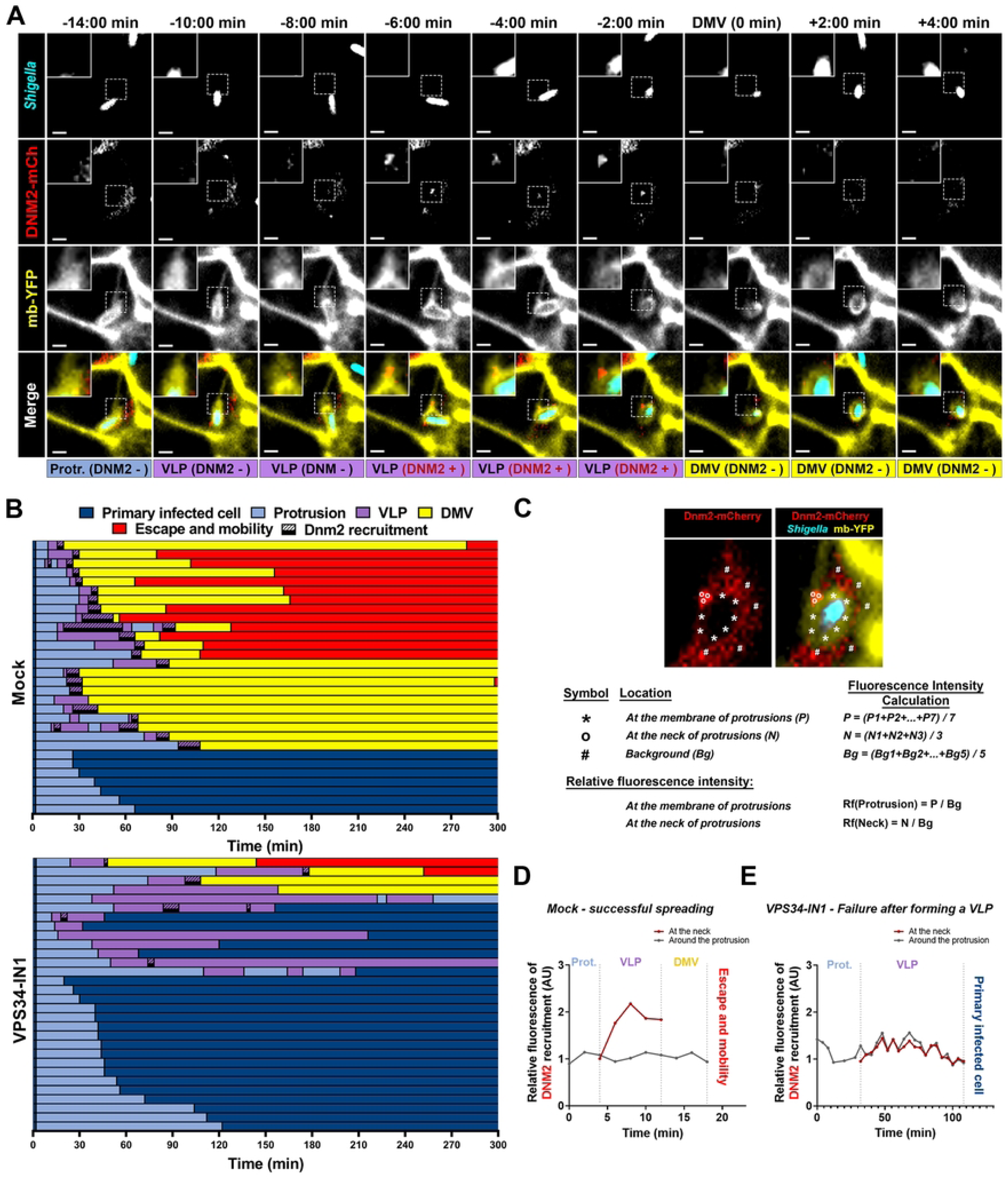
DNM2 promotes *Shigella flexneri* cell-to-cell spread through transient recruitment at the neck of VLPs, leading to DMV formation. **(A)** Representative images of membrane YFP-expressing cells infected with CFP-expressing *S. flexneri* showing the recruitment of DNM2-mCherry during cell-to-cell spread. Scale bar, 2 µm. **(B)** Representative tracking analysis in HT-29 cells infected with CFP-expressing *S. flexneri* in absence (Mock, top panel) or presence of VPS34-IN1 at 500 nM (VPS34-IN1, bottom panel). Each bar represents the tracking of a single bacterium spreading from a non-expressing cell into a DNM2-mCherry expressing cell. At least 30 bacteria were tracked for each condition. Color code: dark blue, primarily infected cells; light blue, protrusions; purple, vacuole-like protrusions (VLPs); yellow, double membrane vacuoles (DMVs) and red, cytosol of adjacent cells; black bar, recruitment of DNM2-mCherry. **(C)** Representative example of relative fluorescence intensity calculation. **(D-E)** Graphs representing the relative fluorescence intensity at the membrane of protrusion/VLP/DMV for an example of successful cell-to-cell spread (D), and failure in VLP (E).

### PIK3C3 contributes to *S. flexneri* cell-to-cell spread *in vivo*

Recent studies have revealed infant rabbits as a new *in vivo* model of shigellosis (Kuehl et al., 2020; Yum *et al*., 2019). The analysis of the colon after intra-rectal infection shared many pathogenesis features with human shigellosis, such as bloody diarrhea, epithelial cell invasion, cell-to-cell spread throughout the mucosa, epithelial fenestration and massive immune cell infiltration (Koseoglu *et al*., 2022; Kuehl *et al*., 2020; Weddle *et al*., 2022; Yum *et al*., 2019). To validate our *in vitro* approach showing an involvement of PIK3C3 in the dissemination of *S. flexneri*, we intra-rectally infected infant rabbits for 6 h with wild type *S. flexneri* strain in presence of the PIK3C3 inhibitor SAR405 and evaluated the capacity of *S. flexneri* to disseminate into the colonic epithelium. Infected colons were immuno-stained with antibodies targeting E-cadherin (green) and *S. flexneri* (red) to respectively visualize epithelial cells and bacteria, using epifluorescence microscopy (Fig 6A). We observed a decrease in the size of the infection foci formed in animals treated with SAR405 compared to mock-treated animals (Fig 6B). We previously showed the inhibition of PIK3C3 by VPS34-IN1 did not impact actin-based mobility at the concentrations used in this study (Fig 1I). To determine whether the high dose of SAR405 used *in vivo* may have an off-target effect on actin-based motility, we infected HT-29 cells in presence of high concentration of SAR405 and determined the percentage of bacteria presenting an actin tail using confocal microscopy. The highest SAR405 concentration tested did not significantly affect the percentage of bacteria with an actin tail, confirming that PIK3C3 inhibition does not impact *S. flexneri* dissemination through actin-based motility (Fig 6C). These results therefore support the notion that PIK3C3 not only supports *S. flexneri* dissemination *in vitro,* but also *in vivo*.

**Fig 6.**
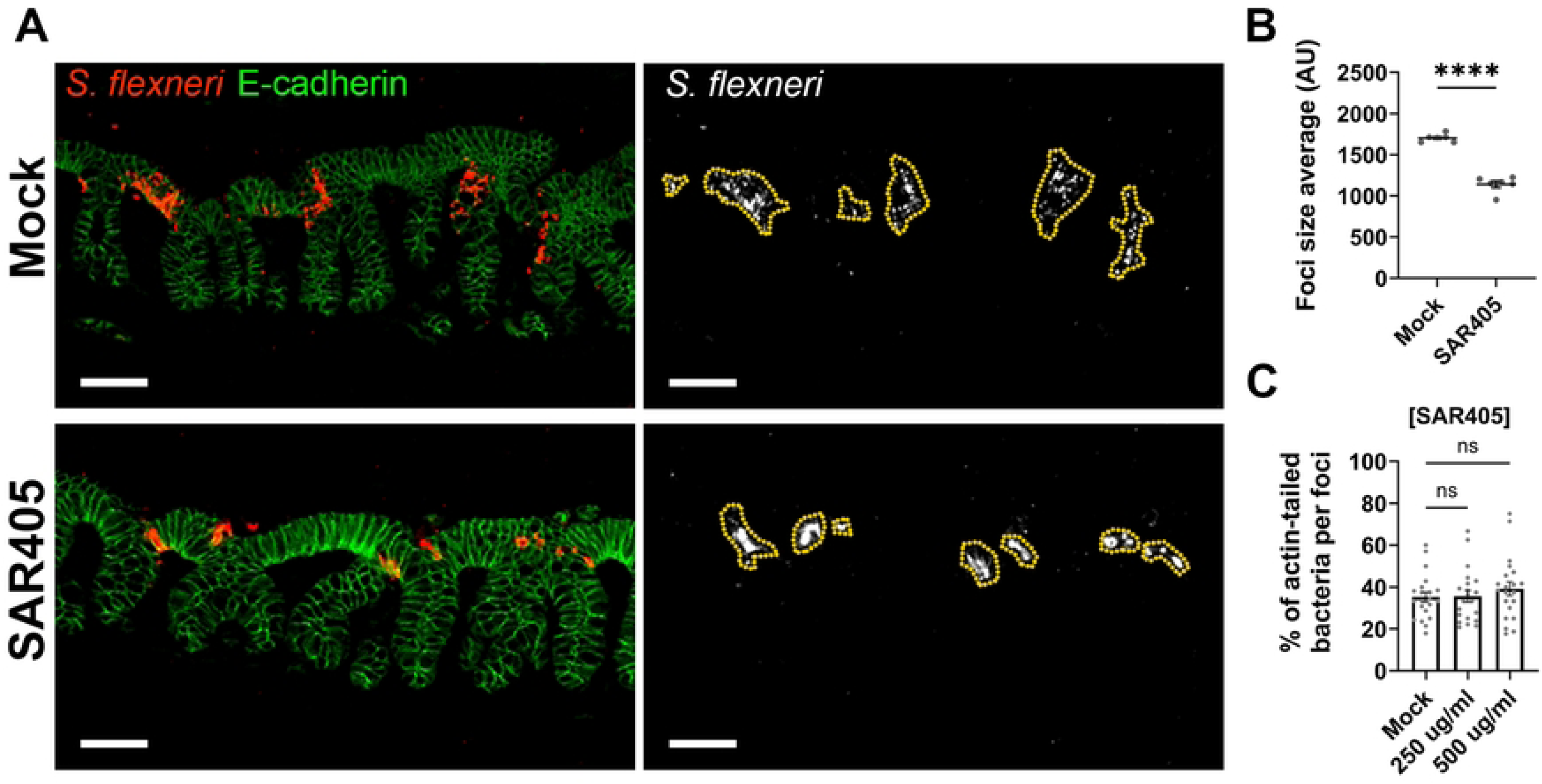
PIK3C3 is required for efficient cell-to-cell spread *in vivo*. **(A)** Representative images of infant rabbit colon sections showing infection foci formed by *S*. *flexneri* (red) in E-cadherin-positive epithelial cells (green) 6 hours post-infection. Scale bar, 50 μm. **(B)** Foci size average (area in arbitrary units) formed by *Shigella* in infant rabbit colon sections in absence and in presence of SAR405. Each dot represents the average foci size per animal. Statistical analysis, unpaired t-tests; ns, not significant; ****, p<0.0001. **(C)** Percentage of bacteria displaying an actin tail per infection focus 6 hours post-infection in presence of SAR405 in HT-29 cells. Eight foci were analyzed per replicate in three independent biological replicates. Statistical analysis, two-way ANOVA with Sidak’s multiple comparisons; ns, not significant.

In summary, we propose a model of *Shigella* dissemination in which the PIK3C3- dependent production of PtdIns(3)P at the VLP stage of cell-to-cell spread supports the resolution of VLP into DMV through recruitment of the DNM2-dependent membrane scission machinery (Fig 7).

**Fig 7.**
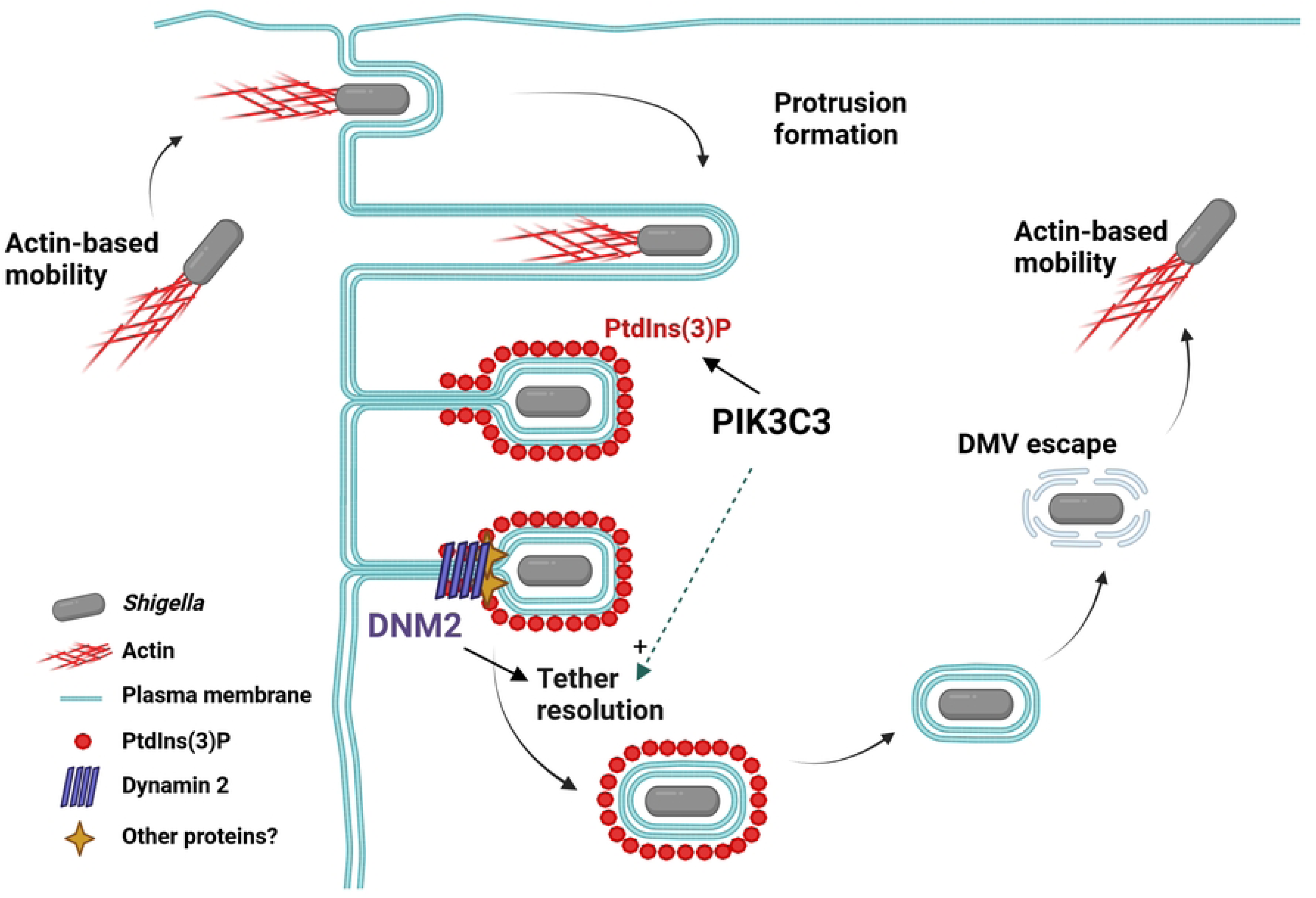
Model of *Shigella* cell-to-cell spread supported PIK3C3 and DNM2. After escape from primary vacuoles and acquisition of actin-based mobility, *S. flexneri* projects into adjacent cells through formation of membrane protrusions. Collapse of the actin cytoskeleton in protrusions leads to the formation of VLPs. PIK3C3-dependent formation of PtdIns(3)P at the protrusion/VLP membrane is required for the recruitment at the VLP neck of the membrane scission machinery, DNM2, leading to the formation of DMVs. *S. flexneri* escape from DMVs does not rely on PIK3C3-dependent formation of PtdIns(3)P. After escaping DMVs, *S. flexneri* resumes actin-based motility in the cytosol of adjacent cells.

## Discussion

### Role of DNM2 in DMV formation

As *S. flexneri* spreads from cell to cell, it forms membrane protrusions that are engulfed by adjacent cells, which leads to the formation of DMVs. Our previous work showed that the resolution of *S. flexneri* protrusions is a multi-step process involving the formation of an intermediate structure termed vacuole-like protrusions (VLPs) with reference to the membranous tether that links VLPs to primary infected cells (Dragoi and Agaisse, 2015). The resolution of the membranous tether leads to the formation of double-membrane vacuoles (DMVs). While the formation of VLPs is supported by signaling events, such as PIK3C2α and tyrosine kinase signaling regulating the collapse of the actin cytoskeleton in protrusions, the mechanisms supporting the resolution of VLPs in DMVs remain unclear. Here, we show that the remodeling of the membranous tether requires the membrane scission machinery, DNM2. DNM2 was previously shown to be important for efficient *S. flexneri* dissemination using genetic ablation and plaque assay as readout of cell-to-cell spread (Fukumatsu *et al*., 2012). Moreover, fluorescence microscopy experiments on fixed samples suggested that DNM2-GFP was recruited to protrusions when expressed in adjacent cells. This is in contrast with our present work showing that DNM2 is recruited to the junction between the membrane surrounding bacteria and the membranous tether, which we refer to as the neck of VLPs. Importantly, the use of time-lapse microscopy allowed us to unambiguously demonstrate that the recruitment of DNM2 coincides with the remodeling of the VLP membranous tether, leading to DMV formation. This is conceptually similar to the mechanisms supporting endosome formation, where DNM2 forms a contractile ring around the membrane of nascent vesicles, leading to membrane scission (Antonny et al., 2016; Morlot et al., 2012). Our work is thus in agreement with the notion that the resolution of VLPs into DMVs is mediated by the membrane scission machinery DNM2 at the neck of VLPs.

### PIK3C3/PtdIns(3)P and DNM2 recruitment to VLPs

Our present work led to the identification of PIK3C3 as a critical regulator of *S. flexneri* dissemination. We showed that PIK3C3 inhibition prevented the formation of PtdIns(3)P in the outer membrane of protrusions, in adjacent cells. Interestingly, PtdIns(3)P was formed either before or slightly after VLP formation, whereas DNM2 recruitment occurred later on, at the time of DMV formation. Moreover, while DNM2 recruitment was localized at the neck of VLPs, PtdIns(3)P was observed in the membrane surrounding bacteria and all along the membranous tether. Since the recruitment of DNM2 was strictly dependent on PIK3C3, these results suggest that PtdIns(3)P formation may act as an upstream signal regulating DNM2 recruitment. Interestingly, in the context of clathrin-mediated endocytosis, DNM2 recruitment requires adaptor proteins, such as SNX9, that display domains that specifically bind PtdIns(3)P (Yarar et al., 2008). Moreover, adaptors such as SNX9 also harbors BAR domains involved in sensing membrane curvature, thereby sensing the temporal coincidence of phosphoinositide signaling, such as PtdIns(3)P, and membrane curvature at the neck of deeply invaginated clathrin-coated pits (Daste et al., 2017; Pylypenko et al., 2007). Altogether, we propose that the combination of PtdIns(3)P formation in the outer membrane of VLPs together with the membrane curvature at the neck of VLPs specify the recruitment of a set of adaptors that in turn recruit DNM2. Further investigations on PtdIns(3)P-binding proteins, such as SNX9, and their potential role will be necessary to better understand the mechanisms supporting PtdIns(3)P formation and DNM2 recruitment to VLPs.

### Differential requirement for PIK3C3 in bacterial dissemination

In addition to *S. flexneri*, we found that PIK3C3 is also required for *L. monocytogenes* spread from cell-to-cell. However, our tracking experiments showed that, unlike *S. flexneri*, PIK3C3 is not required for resolving protrusions into vacuoles during *L. monocytogenes* dissemination, but for escaping DMVs. During the initial entry of *L. monocytogenes*, primary vacuole escape requires acidification of the vacuolar compartment bearing characteristics of late endosome. Acidification is mandatory for activation of the pore-forming toxin listeriolysin O (LLO), which challenges the integrity of the vacuole membrane (Beauregard et al., 1997; Henry et al., 2006; Schnupf and Portnoy, 2007). During endosomal maturation, the progressive replacement of Rab5 by Rab7 is essential for proper acidification of the endosomal compartment through fusion with lysosomes. Interestingly, a recent study showed that PIK3C3 is required for the recruitment of Rab7 to the *L. monocytogenes* primary vacuole and bacterial escape (Cain et al., 2023). Since LLO is also required for escape from DMVs (Gedde et al., 2000), we speculate that PIK3C3 is required for proper acidification of DMVs, thereby facilitating LLO activation and disruption of DMV membrane. Remarkably, *S. flexneri* and *L. monocytogenes* have both co-opted PIK3C3 for facilitating cell-to-cell spread, but through different mechanisms and at different steps during the dissemination process.

### Chemical screens and therapeutic interventions

Our *in vitro* screen for inhibitors of human kinases that would affect *S. flexneri* dissemination led to the identification of VPS34-IN1, a potent inhibitor of PIK3C3. We confirmed the specificity of our findings by showing that an independent PIK3C3 inhibitor, SAR405, also impacted *S. flexneri* dissemination in HT-29 cells. Moreover, we extended this observation *in vivo* by showing that treatment with SAR405 inhibited *S. flexneri* cell-to-cell spread in infant rabbits. These results open the possibility of using treatments that instead of targeting bacterial physiology, would affect host cell processes required for successful infection. Such alternative therapeutic interventions are potentially important in the context of rampant cases of shigellosis involving multi-drug-resistant strains worldwide. Targeting PIK3C3 has already been proposed to improve the treatment of certain types of cancer (Noman et al., 2020; Yu et al., 2024). However, given the role of PIK3C3 in numerous physiological processes, the use of PIK3C3 inhibitors has been abandoned due to safety concerns (Dolgin, 2019; Jaber et al., 2012; Noman *et al*., 2020). Under our experimental conditions, SAR405 was not toxic during the first hours of infection, when we could readily measure the size of the infection foci formed upon bacterial dissemination. Further studies will be required to determine whether long-term treatment with SAR405 would be toxic and thus not suitable for treating pathological outcomes during shigellosis.

## Materials and Methods

### Bacterial strains, mammalian cells and growth conditions

The wild type *Shigella flexneri* strain used in this study is *S. flexneri* 2a str. 2457T (Labrec et al., 1964). To visualize bacteria by fluorescence microscopy, the pMMB207 plasmid conferring chloramphenicol resistance and harboring the gene encoding cyan fluorescent protein (CFP) under the control of an isopropyl-beta-D-thiogalactopyranoside (IPTG)- inducible promoter was introduced by electroporation, resulting in a CFP-expressing *S. flexneri* strain, as previously described (Weddle and Agaisse, 2018b). LB agar plate supplemented with 0.01% Congo Red and LB broth was used to grow *S. flexneri*. For animal experiments, Tryptic soy broth (TSB) was used to culture *S. flexneri*. *Escherichia coli* DH5α strain was used for cloning steps and was grown in LB medium. The wild type *Listeria monocytogenes* strain used in this study is *L. monocytogenes* str. 10403S (Bishop and Hinrichs, 1987).To visualize bacteria by fluorescence microscopy, the pHT315 plasmid conferring erythromycin resistance and harboring the gene encoding CFP or green fluorescent protein (GFP) under the control of an IPTG-inducible promoter was introduced by electroporation, resulting in a CFP- or GFP-expressing *L. monocytogenes* strain as previously described (Talman *et al*., 2014). Brain heart infusion (BHI) agar plate and BHI broth were used to grow *L. monocytogenes*. When appropriate, chloramphenicol (10 µg/mL final concentration), ampicillin (100 µg/mL final concentration) and erythromycin (10 µg/mL final concentration) were added to grow bacteria.

HT-29 cell lines were cultured at 37°C with 5% CO2 in McCoy’s 5A medium (Gibco) supplemented with 10% heat-inactivated fetal bovine serum (HI-FBS; Gibco). The HEK 293T cell line was used to make lentiviruses and was cultured at 37°C with 5% CO2 in DMEM supplemented with 10% HI-FBS. Dulbecco’s Phosphate Buffered Saline (DPBS; Gibco) was used to wash the cells. HT-29 and HEK 293T cells were lifted with Trypsin-EDTA (Gibco) at 0.25% and 0.05%, respectively.

### Antibodies and inhibitors

The following antibodies were used for western blot (WB) or Immunofluorescence (IF): rabbit monoclonal anti-PIK3C3 (Cell signaling technology cat#4263, WB 1:1,000), rabbit polyclonal anti-Actin (Millipore sigma A2066, WB 1:10,000), rabbit polyclonal anti-DNM2 (Abcam ab3457, WB 1:1,000), HRP-conjugated goat anti-rabbit IgG (Jackson, WB 1:10,000), mouse monoclonal anti-E-cadherin (BD Biosciences 610181, IF 1:100), rabbit polyclonal anti-*Shigella sp*. (Virostat cat#0901, IF 1:100), AlexaFluor 594-conjugated goat anti-mouse IgG (Molecular Probes, IF 1:100), AlexaFluor 514-conjugated goat anti-rabbit (Molecular Probes, IF 1:100). For F-actin visualization, AlexaFluor 594-conjugated Phalloidin (Invitrogen A12381, IF 1:1000) was used.

PIK3C3 inhibitors SAR405 (Selleckchem, Cat#S7682) and VPS34-IN1 (MedChemExpress, Cat#HY-12795) were dissolved in 100% dimethyl sulfoxide (DMSO), aliquoted and conserved at -80°C (Bago et al., 2014; Ronan *et al*., 2014).

### Host kinase screen

The kinase inhibitor library was purchased from Selleckchem (L1200). It contained 430 compounds at 10 mM. Compounds were first diluted to 1 mM with DMSO, then diluted to 20 µM with water in 96-well plates, which we refer to as the chemical library test plates. For screening, HT-29 cells were seeded in 96-well plate (Corning, 3904) four days prior to infection, at a final density of 3,000 cells/100 µl/well in McCoy’s Medium supplemented with 10% HI-FBS. The day before infection, *S. flexneri* was grown overnight in 2ml LB at 37°C. The day of infection, each well received 5 ul of the chemical library test plates, thereby achieving a final compound concentration of 1 µM . After incubating the plates at 37°C for 1 hour, 10 µl of McCoy’s medium containing 10% HI- and a 50-fold dilution of *S. flexneri* overnight culture were added to each well. The plates were centrifuged for 5 minutes at 1,000 rpm and incubated at 37°C for 1 hour. After incubation, 50 µl of 10% FBS McCoy’s medium, 40 mM IPTG and 200 µM Gentamicin were added to each well, thereby achieving a final concentration of 10mM IPTG and 50µM Gentamicin. Plates were incubated at 37°C for 7 hours. The medium was then removed from each well and replaced with 50 µl of PBS (Gibco) 4% paraformaldehyde (Electron Microscopy Sciences). Plates were incubated at room temperature for 20 minutes. Fixed cells were washed three times with 100 µl PBS and kept at 4°C until staining and imaging.

### Generation of lentiviruses and infection of HT-29 cells

Lentiviruses were produced from HEK 293T cells to generate HT-29 cell lines stably expressing yellow fluorescence proteins targeted to the plasma membrane (mbYFP), mCherry fused to the PX domain of the p40^phox^ subunit of the superoxide-producing phagocyte NADPH-oxidase (referred as mCherry-PX or mCh-PX), or mCherry fused to DNM2 (referred as DNM2-mCherry or DNM2-mCh). The pMXsIP-mCherry vector was used for cloning. pMX-mbYFP was already constructed and used in previous studies (Weddle and Agaisse, 2018b). pMX-mCherry-PX was cloned after amplification of PX sequence from pMX-YFP-PX (Dragoi and Agaisse, 2015) using the following primers: 5PX_EcoRI (5’ GAAGAATTCCATGGCTGTGGCCCAGCAGCTG) and 3PX_NotI (5’ GCGGCGGCCGCTTATTGGCGGAGTGCCTGGGGCACC) and inserted in pMXsIP-mCherry after enzymatic digestion with EcoRI and NotI and ligation. pMX-DNM2-mCherry was cloned by insertion of Dyn2-mCherry fragment into pMXsiP-mCherry successively digested with PacI (NEB), blunted with DNA polymerase I, Large (Klenow) fragment and digested with NotI (NEB). Insert containing DNM2-mCherry was extracted from pDyn2- mCherry N1 (Addgene, Cat#27689) after digestion with BglII (NEB), blunted with DNA polymerase I, Large (Klenow) fragment (NEB, cat# M0210S), and digested with NotI (NEB). To make lentiviruses, pMX- vectors were co-transfected in HEK 293T cells with packaging vectors pCMVΔ8.2Δvpr (HIV helix packaging system) and pMD2.G (a vesicular stomatitis virus glycoprotein) using X-tremeGENE HP DNA transfectant (Roche) (Kissler et al., 2006). After two days, supernatants containing lentiviruses were collected and filtered through 0.45 µm syringe filters to remove HEK 293T cells. HT-29 cells were infected by lentiviruses for one day and puromycin at 5 µg/mL was used to select mb- YFP, DNM2-mCherry or mCherry-PX expressing HT-29 cells. Puromycin was removed after obtaining the corresponding stable cell lines.

### Treatment with siRNA duplexes

All siRNA duplexes used in this study were purchased from Dharmacon (Horizon discovery). For PIK3C3 knockdown, siRNA #A (cat# D-005250-01) and siRNA #B (cat# D-005250-04) were used. For DNM2 knockdown, siRNA #A (cat# D-004007-02) and siRNA #B (cat# D-004007-18) were used. N-wasp siRNA (cat# D-006444-02) was used as positive control for *S.flexneri* cell-to-cell spread defect. siRNA buffer alone was used as negative control (mock). Four days before infection, 50nM of siRNA was transfected in HT-29 cells by reverse-transfection using DharmaFECT 1 (Horizon discovery) following the manufacturer’s protocol. Knockdown efficiency was verified by western blot on cells collected on the day of infection.

### Infection of HT-29 cells and foci size analysis

Cell-to-cell spread was determined by infecting a 4-day old confluent monolayer of HT-29 cells in 96-well plates (Corning, cat#3904) using exponential phase CFP-expressing *S. flexneri*, or overnight culture of GFP-expressing *L. monocytogenes* as previously described (Koseoglu *et al*., 2022; Weddle *et al*., 2022). Briefly, bacteria were diluted in cell media and added directly to the cell medium. Plate were centrifuged at 1,000 rpm for 5 min at room temperature to initiate and synchronize infection. After one hour of incubation at 37°C in 5% CO2 to allow for bacterial invasion, IPTG (10 mM final concentration) and gentamicin (50 μg/ml final concentration) were added to wells to induce the expression of CFP (*Shigella*) or GFP (*Listeria*) and to kill extracellular bacteria, respectively. The plate was incubated for 7 h or 15 h at 37°C in 5% CO2. Infection was stopped after fixation using 4% paraformaldehyde (Electron Microscopy Sciences, cat# 15710) in PBS. The integrity of the cell monolayer was verified using Hoechst staining. PIK3C3 inhibitors VPS34-IN1 (MedChemExpress, Cat#HY-12795) and SAR405 (Selleckchem, Cat#S7682) were added after 1 hour of infection at same time as IPTG and Gentamicin, and DMSO (0.1 % final concentration) was used as mock condition. After fixation, plates were imaged using an ImageXpress Micro imaging system (Molecular Devices). The image analysis was performed with the ImageXpress imaging software (Molecular Devices), as previously described (Dragoi and Agaisse, 2014a). Image analyses were conducted on at least 50 bacterial infection foci in each condition of each biological replicate.

### Confocal microscopy on fixed samples

To determine the effect of PIK3C3 inhibitors on Actin-tail formation or on cell-to-cell spread, mbYFP-expressing HT-29 cells were cultured in McCoy’s medium on round cover slips, German glass 12 mm (Thomas Scientific) in 24-well plate (Costar, Cat#3526) for four days. Medium was replaced at each stage of the infection process by fresh medium containing bacteria or IPTG/Gentamicin. Infection was stopped by adding PBS containing 4% PFA at the time points indicated in the legend of the corresponding figure. Coverslips were stained using AlexaFluor 594-conjugated Phalloidin (Invitrogen A12381, IF 1:1,000) to visualize F-actin. DABCO was used as mounting solution. Coverslips were imaged using a Leica DMI 8 spinning-disc confocal microscope driven by the iQ software (Andor). Images were analyzed using the Imaris software (BitPlane).

### Live confocal microscopy

Bacterial cell-to-cell spread was monitored using time-lapse confocal microscopy. HT-29 cells were seeded in collagen-coated (Sigma- Aldrich, cat#C3867, 1:100) eight-well chambers Lab-Tek II (Thermo Scientific, cat#155409) at 37°C in 5% CO2. A mixed population of HT-29 mbYFP and HT-29 mbYFP mCherry-PX or DNM2-mCherry (ratio 1:1) was seeded to visualize PtdIns(3)P formation and DNM2 recruitment during cell-to-cell spread. After four days, the cell monolayer was infected with CFP-expressing *S. flexneri* or CFP-expressing *L. monocytogenes*. After one hour of incubation at 37°C in 5% CO2 to allow for bacterial invasion, IPTG (10 mM final concentration), gentamicin (50 μg/ml final concentration) and VPS34-IN1 (500 nM final concentration) (or DMSO 0.1%) were added to wells to induce CFP expression, kill extracellular bacteria and inhibit PIK3C3 (or mock control, DMSO). Cells were imaged every 2 min for 7 hours with a Leica DMI 8 spinning-disc confocal microscope driven by the iQ software (Andor). Imaris software (BitPlane) was used to generate and visualize movies for analysis. At least 30 bacteria were tracked for each biological replicate per condition. The relative fluorescence intensities for PX and DNM2 recruitment were calculated from intensity values computed by the Imaris Software (BitPlane) as described in Figures 4C and 5C. The mean of fluorescence intensities was calculated from seven sites at the membrane of protrusions (white stars), five sites in the background (white sharp symbols) or three sites at the neck of protrusions (white circle symbols).

### Western blot

Cells were washed with DPBS (Gibco) and harvested directly from the well in 2x Laemmli Buffer supplemented with 10mM DTT. Lysates were boiled for 5 minutes and separated by SDS-PAGE. Proteins were transferred on Amersham™ Protran™ 0.2 µm nitrocellulose blotting membrane (Cytiva). The membrane was incubated for 1 hour at room temperature in blocking buffer (5% skim milk in 1X TBS with 0.05% Tween) before overnight incubation at 4 °C with gentle shaking in presence of primary antibody diluted in blocking buffer. Following three washing steps, the membrane was incubated in blocking buffer with the secondary antibodies (horse radish peroxidase conjugated). Signal was visualized using SuperSignal West Pico PLUS Chemiluminescent Substrate (ThermoFisher Scientific, cat#34577). Images were taken using the Bio-Rad ChemiDoc MP Imaging System. Western blot quantification was performed using ImageJ.

### *In vivo S. flexneri* infection

Pregnant New Zealand White rabbits were purchased from a commercial breeding company (Charles River). Animal studies were conducted using infant rabbits, as previously described (Koseoglu *et al*., 2022; Weddle *et al*., 2022; Yum *et al*., 2019). Briefly, 10-15-day old New Zealand White rabbits were isolated after birth and kept at 30°C, with daily feedings from tranquilized does. *Shigella flexneri* was grown overnight at 37°C in 5 mL of TSB per animal. Bacterial culture was pelleted and resuspended in 200 µL PBS containing SAR405 at 2 mg/mL or 2.5% DMSO. Kits were anesthetized with 5% isoflurane and rectally inoculated with 200 μL of bacterial suspension.

University of Virginia Institutional Biosafety Committee and the Institutional Animal Care and Use Committee reviewed and approved animal protocol #4161. The care of the does and infant rabbits was executed according to standard operating procedures developed in coordination with the veterinary and animal care staff of the Center for Comparative Medicine at the University of Virginia.

### Colon collection and Histology

After 6 hours of infection, infected infant rabbits were euthanized by CO2 inhalation and euthasol injection and the distal colon was harvested. Infected colons were rinsed in PBS and flushed with modified Bouin’s fixative for paraffin sections. Samples were cut open longitudinally and displayed in cassettes as swiss-rolls. Cassettes were immersed in neutral-buffered formalin for 48 hours. Tissues were preserved in 70% Ethanol before loading onto a tissue processor for dehydration and paraffin infiltration. After manual embedding into a paraffin block, paraffin sections were cut at 5 μm on a Leica microtome. Paraffin sections were stained with hematoxylin and eosin at the research histology core facility at University of Virginia School of Medicine. Colons were imaged using ScanScope Slide Scanner (Leica Biosystems). Epithelial fenestration was measured along the entire colon using the Aperio software and % fenestration (length of colon with fenestrated epithelium/total length of colon x 100) was calculated for each colon.

### Immunofluorescence on colon sections

Colon paraffin sections were sequentially processed via deparaffinization, and re- hydration, as previously described (Yum *et al*., 2019). Antigen retrieval in pre-heated citric acid-based buffer (1:100 in water, Vector Laboratories, Cat#H-3300) was performed in a pressure cooker (Instant Pot) for one minute on the high-pressure setting, followed by a “quick release”. Slides were rinsed three times with PBS, permeabilized for 10 min at room temperature with PBS-Triton 0.1% and blocked for one hour at room temperature in PBS containing 5% bovine serum albumin (BSA) and 2% normal goat serum (NGS). Primary antibodies incubation was conducted at 4°C overnight in a humidified chamber with rabbit anti-*Shigella sp.* antibody (Virostat, 1:100) and mouse anti-E-cadherin antibody (BD Biosciences, 1:100) diluted in PBS containing 5% BSA and 2% NGS. Slides were washed three times with PBS and incubated with AlexaFluor 594-conjugated goat anti-mouse IgG (Molecular Probes, IF 1:100), AlexaFluor 514-conjugated goat anti-rabbit (Molecular Probes, IF 1:100) in PBS containing 5% BSA and 2% NGS for 2 h at room temperature. Coverslips were mounted with ProLong Gold Antifade Mountant (Thermo Fisher). Entire colons were imaged using a Nikon TE2000 microscope equipped for multi- color imaging including motorized stage and filter wheels and a Hamamatsu Orca ER Digital CCD Camera. The size of the *Shigella* infection foci was determined using the Region function of the MetaMorph software (Molecular Devices).

## Statistical analysis

Statistical analyses of the values were conducted using the GraphPad Prism 9.0.2 software. Analysis of a single variable across two groups was performed with unpaired t- test to determine statistical significance. Experiments with multiple groups were analyzed with one-way ANOVA followed by Tukey’s multiple comparisons. P-values were represented on figures as follows: ns, not significant; *, p<0.05; **, p<0.005; ****, p<0.0001.

## Acknowledgments

We thank members of the Agaisse lab at the University of Virginia for discussions and comments on the manuscript. We thank Valerie Spencer and Cierra Roach for technical assistance with animal work. We thank Hannah Collier, Alice Kweon, Zack Lifschin, Liam Myers, and Ellie Wood for technical assistance with infant rabbit nursing. We thank Sheri Vanhoose and the Research Histology Core for preparing paraffin sections.

## Author contributions

**Steven J Rolland**

**Roles:** Conceptualization, Investigation, Methodology, Formal Analysis, Validation, Visualization, Writing – Original Draft Preparation

**Affiliation:** Department of Microbiology, Immunology, and Cancer Biology, University of Virginia School of Medicine, Charlottesville, Virginia, United States of America.

**Zachary J Lifschin**

**Roles:** Investigation, Methodology, Formal Analysis

**Affiliation:** Department of Microbiology, Immunology, and Cancer Biology, University of Virginia School of Medicine, Charlottesville, Virginia, United States of America.

**Erin A Weddle**

**Roles:** Investigation, Methodology, Formal Analysis

**Affiliation:** Department of Microbiology, Immunology, and Cancer Biology, University of Virginia School of Medicine, Charlottesville, Virginia, United States of America.

**Lauren K Yum**

**Roles:** Investigation, Methodology, Formal Analysis

**Affiliation:** Department of Microbiology, Immunology, and Cancer Biology, University of Virginia School of Medicine, Charlottesville, Virginia, United States of America.

**Tsuyoshi Miyake**

**Roles:** Investigation, Methodology

**Affiliation:** Department of Microbiology, Immunology, and Cancer Biology, University of Virginia School of Medicine, Charlottesville, Virginia, United States of America.

**Daniel A Engel**

**Roles:** Conceptualization, Funding Acquisition, Methodology, Project Administration, Supervision, Writing – Review & Editing

**Affiliation:** Department of Microbiology, Immunology, and Cancer Biology, University of Virginia School of Medicine, Charlottesville, Virginia, United States of America.

**Hervé F Agaisse**

**Roles:** Conceptualization, Funding Acquisition, Methodology, Project Administration, Supervision, Writing – Review & Editing

**Affiliation:** Department of Microbiology, Immunology, and Cancer Biology, University of Virginia School of Medicine, Charlottesville, Virginia, United States of America.

## Supporting information

**S1 Table. Host kinase screening results.**

Hits obtained in three biological replicates (R1-3) showing infection foci size and corresponding Z-Scores. Hits were selected on the basis of a decrease of infection foci size by at least 2 standard deviations (stdev WT), i.e. Z-Score < -2, with respect to the average (avg) foci size in control wells (avg WT) containing DMSO only.

**S1 Fig. *Shigella flexneri* tracking analysis. (A-B)** Representative tracking analysis of second (A) and third (B) biological replicates using live-fluorescence confocal microscopy of CFP-expressing *S. flexneri* during cell-to-cell spread in the absence (left panel) or presence (right panel) of VPS34-IN1 (500 nM). Each bar represents the tracking of a single bacterium over 4h30. Color code: dark blue, primarily infected cells; light blue, protrusion; purple, vacuole-like protrusion (VLP); yellow, double membrane vacuole (DMV) and red, escape and actin-based motility in the cytosol of adjacent cells.

**S2 Fig. Example of cell-to-cell spreading failure in VLP upon VPS34-IN1 treatment.** Live-fluorescence confocal microscopy images of membrane YFP-expressing HT-29 cells infected with CFP-expressing *S. flexneri* in presence of VPS34-IN1 (500 nM). The tracked bacterium forms a protrusion that successfully resolves into a VLP, but does not resolve into a DMV. The VLP subsequently retracts back to the primary infected cell. The arrows indicate the membranous tether formed during the VLP stage. Scale bar, 2 μm.

**S3 Fig. PtdIns(3)P is produced at DMV membrane, but not VLP membrane, during *Listeria* cell-to-cell spread.** Live-fluorescence confocal microscopy images of membrane YFP-expressing HT-29 cells infected with CFP-expressing *L. monocytogenes* spreading from a mCherry-PX non-expressing cell into a mCherry-PX expressing cell. Unlike *S. flexneri* that forms VPLs, the frame before DMV formation shows a protrusion in cells infected with *L. monocytogenes*. Scale bar, 2 μm.

**S4 Fig. *Listeria monocytogenes* tracking analysis. (A-B)** Representative tracking analysis of the second (A) and third (B) biological replicates using live-fluorescence confocal microscopy of CFP-expressing *L. monocytogenes* during cell-to-cell spread in absence (left panel) or presence (right panel) of VPS34-IN1 (500 nM). Each bar represents the tracking of a single bacterium over 4h. Color code: dark blue, primarily infected cells; light blue; yellow, double membrane vacuole (DMV) and red, escape and mobility in the cytosol of adjacent cells.

**S5 Fig. DNM2 promotes *Shigella flexneri* cell-to-cell spread. (A)** Representative images showing infection foci formed in HT-29 cells treated with two siRNA duplexes targeting DNM2 16 hours post-infection. Scale bar, 100 μm. **(B)** Quantification of foci size (area) as shown in (A). Three independent biological replicates were performed and at least 50 infection foci were analyzed per condition. Each dot represents the average of one independent experiment and error bars represent the standard deviation of the mean. Statistical analysis, one-way ANOVA with Dunnett’s multiple comparisons test; ns, not significant; *, p<0.05; **, p<0.005; ***, p<0.001; ****, p<0.0001. **(C)** Western blot showing knockdown efficiency of two siRNA duplexes targeting DNM2. **(D)** Quantification of the knockdown efficiency in three biological replicates. DNM2 signals were normalized to the corresponding actin signals and knockdown efficiency of siRNA duplexes was calculated relative to mock treated cells. Statistical analysis, one-way ANOVA with Dunnett’s multiple comparisons test; ns, not significant; ****, p<0.0001.

